# Evaluating the ileal and cecal microbiota composition of a 1940 heritage genetic line and a 2016 commercial line of white leghorns fed representative diets from 1940 and 2016

**DOI:** 10.1101/2023.06.08.544207

**Authors:** D. C. Wall, Dana Dittoe, Ramon D. Mahlerios, K. E. Anderson, N. Anthony

## Abstract

This study was conducted to identify and evaluate the differences between the microbiome composition of the ileum and ceca of 1940 and 2016 genetic strains of white leghorns fed representative contemporary diets from those times. Ileal and cecal samples were aseptically collected from both genetic lines at 69 weeks of age. The genomic DNA of the ileal and cecal contents were extracted and the V4 region of the 16S rDNA was sequenced on an Illumina Miseq. Microbiota data were filtered and aligned using the QIIME2 2020.2 pipeline. Alpha and beta diversity metrics were generated and the Analysis of Composition of Microbiomes (ANCOM) was utilized to determine significantly different taxa. Data were considered significant at P ≤ 0.05 for main effects and Q ≤ 0.05 for pairwise differences. Alpha diversity of the ileum and ceca were significantly different (P = 0.001; Q = 0.001; however, no differences between genetic lineage were observed (P > 0.05; Q > 0.05). The beta diversity between the ileum and ceca, as well as between the genetic lines (1940 vs. 2016) were significantly different from one another (P = 0.001; Q = 0.001). Using ANCOM, *Proteobacteria* and *Actinobacteriota* were significantly different than other phyla (P ˂ 0.05) with a higher relative abundance of *Proteobacteria* being observed among treatment groups 2 and 3, while *Actinobacteriota* had higher relative abundance in treatment groups 1 and 4. Among the significantly different genera in the ileum, *Pseudomonas*, *Rhizobiaceae*, *Leuconostoc*, and *Aeriscardovia* were different (P ˂ 0.05) with treatment groups 1 and 4 having a higher relative abundance of *Aeriscardovia*, while treatment groups 2 and 3 had higher relative abundance in both *Pseudomonas* and *Leuconostoc*. In the ceca, *Proteobacteria*, *Firmicutes*, *Actinobacteriota*, and *Euryarchaeota* were significantly different phyla (P ˂ 0.05) with *Firmicutes* having the highest relative abundance across all treatment groups. Among the significantly different genera (*Pseudomonas*, *Leuconostoc*, *Alloprevotella*, and *Aeriscardovia*), *Alloprevotella* had the highest relative abundance across all treatment groups 1 and 2, while *Leuconostoc* and *Pseudomonas* had the highest relative abundance in treatment group 4. Results from this study suggest that genetic makeup in conjunction with the nutritional composition of laying hens influences the cecal and ileal microbiota of corresponding hens.

## Introduction

Diet directly impacts and shapes the composition and subsequent functions encoded by the microbiota within the gastrointestinal tract of poultry (Hubert, S. et al, 2019). The integrity of the gastrointestinal tract (GIT) and the gut microbial community play vital roles in nutrient absorption, development of immunity, and disease resistance (Shang, Y et al., 2018). As such, the microbiota is in constant interaction with the host and has been shown to influence several major functions such as the immunological, physiological, and nutritional status of birds. In addition, poultry is an ideal model for host-microbiota interactions as the microbiota establishment of commercial poultry is relatively independent of that of the parent stock (Zhao L et al., 2013; Rychlik, 2020). Research conducted by Kers et al. (2018) demonstrated differences in the intestinal microbiota composition. Within the same study having controlled conditions they showed that genetics plays a factor that could influence microbiota composition.

The microbiota of the small intestine has been demonstrated to have relatively low diversity. It is estimated that 50% of the total ileal microbiota can be formed by one to five genera only (Rychlik, 2020). Although the ileum may have a relatively smaller abundance of microbiota, the microbiota present within directly impacts ileal digestibility (Ptak, A et al., 2015). Unlike the ileum, the ceca possess the highest microbial richness without poultry and are mainly colonized by anaerobic and microaerophilic microorganisms. Therefore, fermentation of nutrients indigestible to the host occurs with the ceca due to the longer retention time of feed within the ceca (Clench, M. et al., 1995). Compared to the ileum, the cecum microbiota is more diverse and more stable.

A large and diverse microbiota exists within the ceca of laying hens, prolonging the retention time of digesta to approximately 12–20 h., a process known as microbial-based metabolism (Huang, C. B., et al., 2019). It has been estimated that over 10 Log10 CFU per gram of digesta and approximately 100 different taxa exist within the ceca of poultry (Oakley BB, et al. 2014; Kogut, Michael, 2018). These taxa belong to the two major phyla; Gram-positive *Firmicutes* and Gram-negative *Bacteroidetes*, followed by two major phyla: *Actinobacteria* (Gram-positive) and *Proteobacteria* (Gram-negative) (Oakley BB, et al. 2014; Kogut, Michael, 2018). *Firmicutes* and *Bacteroidetes* are generally equally distributed in the cecal microbiota of healthy adult hens comprising 45% of the total microbiota (Schreuder, J., et al., 2020); whereas the abundance of *Actinobacteria* and *Proteobacteria* is only 2–3% of total microbiota (Rychlik, 2020).

Due to the rapid development of DNA sequencing technologies, the composition of the gut microbiota within poultry is rapidly being described. However, there is little understanding of how the microbiota has been altered with the genetic selection pressures utilized in the poultry industry to enhance poultry health and performance. Additionally, there is limited knowledge on how improved diet formulation has altered the microbiota of poultry. Therefore, the aim of the current study was to describe and identify potential differences of the ileal and cecal microbiota composition of two genetic strains of white leghorns; a heritage line from 1940 and a commercial line from 2016 fed diets representative of 1940 and 2016.

## Materials and Methods

### Bird Management and Diet

At 16 weeks of age, a total of 320 white leghorn laying hens (WL40 and WL36) were transported and housed in conventional cages at a laying facility at the North Carolina Chicken Education Unit in Raleigh, NC. The rearing of these birds was carried out in accordance with the Institutional Animal and Use Committee at North Carolina State University (NCSU IACUC) [30]. All birds were randomly divided into 4 treatment groups with 80 hens per treatment N = 320; n = 80; k = 4). Treatment groups were as follows: 1). 2016 hen on 1940 diet; 2). 2016 hen on 2016 diet; 3). 1940 hen on 1940 diet; and 4). 1940 hen on 2016 diet. There were 2 hens per cage consisting of 8 replicate cages. Feed and water were provided ad libitum throughout the experimental period of 69 weeks (**Table 1**). Feed intake and body weight gain were measured on a 28d period resulting in 12 cycles. All animal management and sampling procedures were in accordance with the NCSU IACUC [30].

**Table 1.** Feed Ingredients and Mash. ^1^Diet Compositions

| **Ingredients** | **2016 Layer Diet<sup>2</sup>** | **1940 Layer Diet<sup>2</sup>** |
| --- | --- | --- |
| **Corn** | 940.5 | 1146.38 |
| **Soybean Meal** | 718.0 | 232.57 |
| **Alfalfa Meal** |  | 305.97 |
| **Limestone, gr.** | 145.5 | 124.2 |
| **Coarse limestone** | 50.0 |  |
| **Fat** | 110.0 |  |
| **Phosphate Mono/D** | 17.6 |  |
| **Salt** | 6.8 | 5.0 |
| **D.L. Methionine** | 2.9 |  |
| **T-Premix** | 1.0 |  |
| **Sodium Bi-carb** | 2.0 |  |
| **Prop Acid 505** | 1.0 |  |
| **Choline CL 60%** | 1.3 | 4.0 |
| **Hy-D 62.5 mg/lb** |  |  |
| **Trace Min PMX<sup>3</sup>** | 1.0 |  |
| **L-Vitamin PMX<sup>4</sup>** | 1.0 |  |
| **.06% Selenium<sup>5</sup>** | 1.0 |  |
| **Ronozyme HI P (GT)** | 0.4 |  |
| **Total** | 2000.0 | 2000.0 |
| **Calculated Analysis** |  |  |
| **Protein %** | 20.8 | 20.0 |
| **ME kcal/kg** | 2926 | 1330 |
| **Calcium %** | 4.10 | 0.90 |
| **A. Phos %** | 0.45 | 0.42 |
| **Lysine %** | 1.20 | 0.82 |
| **TSAA %** | 0.81 |  |

<sup>1</sup>Diets were acquired from the North Carolina State University Feed Mill in mash form
<sup>2</sup>Lay diet fed starting no later than 17 weeks
<sup>3</sup>Mineral premix provides per kg of diet: manganese, 120 mg; zinc, 120 mg; iron, 80 mg; copper, 10 mg; iodine, 2.5 mg; and cobalt
<sup>4</sup>Vitamin premix provides per kg of diet: 13,200 IU vitamin A, 4000 IU vitamin D3, 33 IU vitamin E, 0.02 mg vitamin B12, 0.13 mg biotin, 2 mg menadione (K3), 2 mg thiamine, 6.6 mg riboflavin, 11 mg d-pantothenic acid, 4 mg vitamin B6, 55 mg niacin, and 1.1 mg folic acid. 6 Selenium premix = 1 mg
<sup>5</sup>Selenium premix provides 0.2 mg Se (as Na<sub>2</sub>SeO<sub>3</sub>) per kg of diet

### Ileum and Cecal Sample Collection

On the last day of the trial, at 69 weeks of age, 12 hens from each treatment group were randomly selected and euthanized by cervical dislocation. Both the ileum and cecal digesta contents were aseptically collected and stored at -80°C, until further analysis.

#### Microbiome Analysis: 16S rDNA Sequencing

Ileal and cecal digesta samples were shipped at ambient temperature to the University of Arkansas (˂ 72 hrs) for further analysis. The genomic DNA from both ileum and ceca (200mg) were extracted according to the manufacturer’s recommendations using a Qiagen Stoll Mini Kit (Qiagen, Hilden, Germany). Genomic DNA was eluted and quantitated using a NanoDrop 1000 (Thermo Scientific; Waltham, MA, USA). Genomic DNA was diluted to 10ng/µL in Buffer AE (Qiagen, Hilden, Germany). Using custom primers described by Kozich et al., [31] Using a high-fidelity polymerase, Accuprime *pfx* (Invitrogen, Waltham, MA, USA), the V34, V4, and V45 region of 16S rDNA was amplified in a Mastercycler® X50 thermal cycler according to the manufacturer recommendations (Eppendorf, Hamburg, Germany). Using gel electrophoresis, amplification was confirmed, and individual samples were normalized using the SequalPrep™ normalization kit (Applied Biosystems™, Waltham, MA, USA). After samples were in equal molar concentrations, the library was pooled using 5 µL of each sample. The final library size was confirmed using a Qubit fluorometer (Invitrogen, Waltham, MA, USA) and a KAPA qPCR library quantification kit (Roche Sequencing, Pleasanton, CA, USA). Confirmation of the amplified region size (bp) was confirmed using an Agilent 2200 TapeStation (Aglient, Santa Clara, CA, USA).

The library and PhiX control v3 (Illumina, Carlsbad, CA, USA) were diluted to 20nM in HT1 Buffer and denatured in 0.2 N NaOH to generate a final concentration of 12 pM. The diluted library was combined with the subsequent PhiX control v3 (20%, v/v) and loaded onto a MiSeq v2 (500 cycles) reagent cartridge (Illumina, Carlsbad, CA, USA). The resulting sequences were uploaded to BaseSpace (Illumina, San Diego, CA, USA), NCBI Sequence Read Archive (Project Accession), and Github (Lab repository).

### Statistical and Bioinformatics Analysis

The QIIME2 pipeline (version 2020.11) was utilized for sequencing data processing (Bolyen, E. et al., 2019). Demultiplexed reads were downloaded from the Illumina BaseSpace website and were uploaded into Qiime2-2020.11 using Casava 1.8 paired-end demultiplexed format (via Qiime tools import). Demultiplexed reads were denoised and filtered in DADA2 via q2-dada2 (Callahan, B. J., et al, 2016). Using mafft, the observational taxonomic units (OUT’s) were aligned and a rooted phylogenetic tree was generated with fastree2 via q2-phylogeny (Price, M. N et al., 2010). Punitive OTUs were identified against SILVA (Silva 138 99% OTUs full-length sequences) (Quast, C., et al., 2013; Yilmaz, P., et al., 2014; Glöckner, F. O., et al., 2017) using the sk-learn Bayesian algorithm (via q2-feature-classifier) (Bokulich, N. A., et al., 2018).

Alpha and Beta diversity were analyzed using core metrics results in QIIME2. Alpha diversity was determined using the Shannon diversity index. Observed OTUs, Pielou’s Evenness, and Faith’s phylogenetic diversity, which included the Kruskal-Wallis tests for pairwise differences (Kruskal, W., et al., 1952). Beta diversity metrics were analyzed with qualitative and quantitative indices, Bray Curtis, Jaccard, and Unweighted and Weighted UniFrac Distance Matrix (Lozupone, C., et al., 2005; Lozupone, C. A., et al., 2007). PERMANOVA, a multivariate form of ANOVA with permutations to reduce bias, was used to determine if the distribution and abundances among beta diversity metrics were different. Interactions between the organ site and treatment were identified using ANOVA (via q2-longitudinal (Bokulich, N.A., et al., 2018) and ADONIS (Anderson, M.J. 2001) for Alpha and Beta diversity metrics. Pairwise differences for all diversity metrics (Kruskal Wallis and PERMANOVA) were considered significant when P ≤ 0.05 and Q ≤ 0.05. The Q values were used as they include the False Discovery Rate, which accounts for the occurrence of type I errors, rejecting a true null hypothesis (false positive) when conducting multiple comparisons. Lastly, ANCOM (analysis of the composition of microbiomes) was conducted according to Mandal et al. (2015) to determine differentially abundant taxa.

## Results

### Alpha Diversity

There were 4 metrics used to evaluate alpha diversity. The Species diversity or Shannon index determined how diverse or how different the species were within an ecosystem. Species richness or OTU count determines how many different and unique OTUs can be detected in a microbial ecosystem within a sample. Pielou’s evenness determined how evenly the species were distributed within an ecosystem. Faith’s Phylogenetic Diversity was used to determine diversity within the ecosystem using the sum of all the phylogenetic branch lengths of the OTUs in the ecosystem. Among all the diversity indices, significant differences were observed between the microbiota of both the ceca and ileum (P = 0.001; Q = 0.001; **Figure 1**) however, no differences were observed between the four treatment groups (P > 0.05; Q > 0.05; **Supplemental Table 1**).

**Figure 1.**
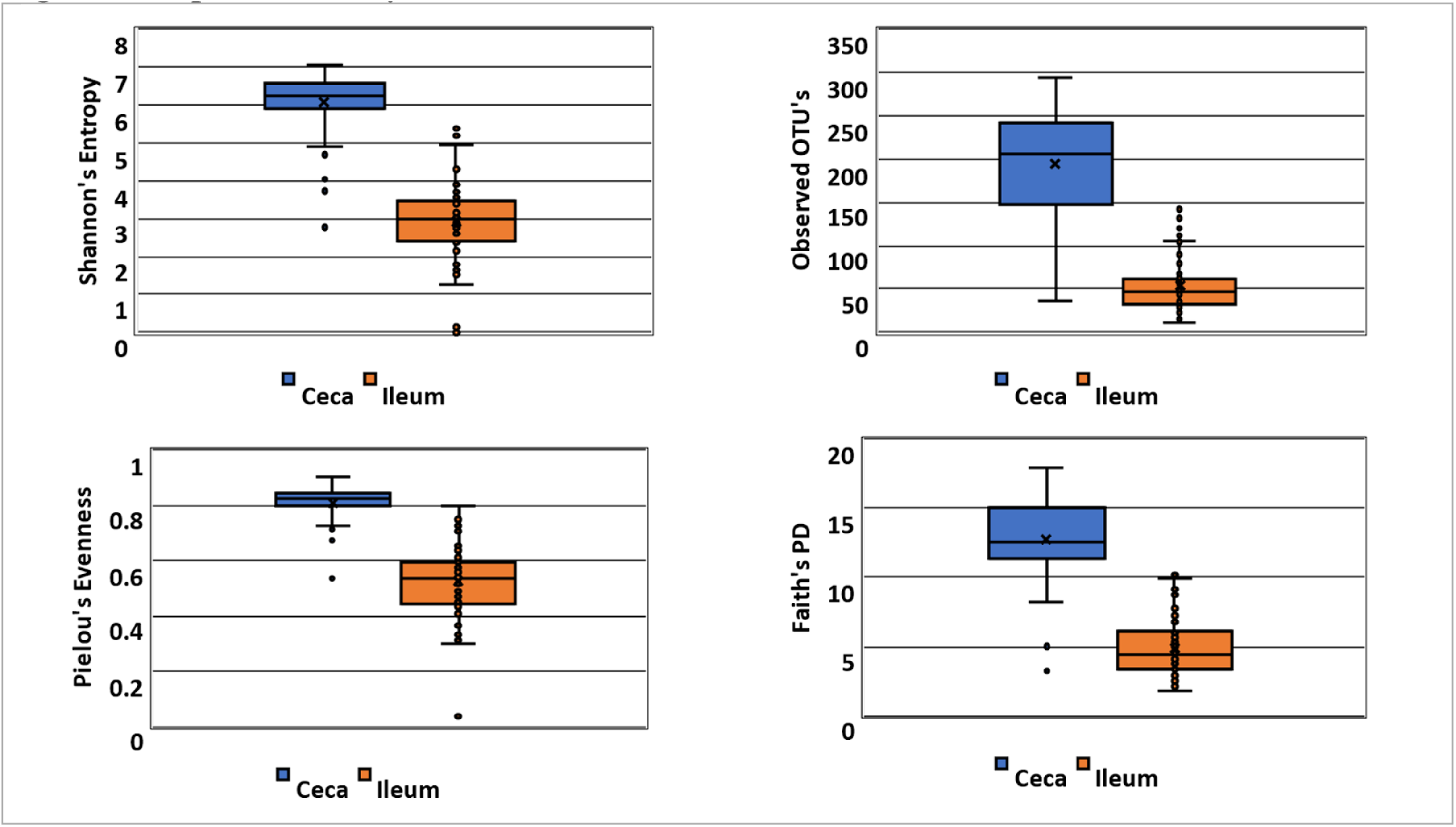
Alpha Diversity of Ceca and Ileum. ^1^Alpha diversity indices of ileum and ceca

### Beta Diversity

There were 4 metrics used to evaluate beta diversity. Bray-Curtis dissimilarity compared the microbial ecosystems to one another based on the differences in microbial abundances between two samples on a scale from 0 to 1 with 1 meaning completely different species abundances. Jaccard distance compared the microbial ecosystems to one another based on the presence or absence of species on a 0 to 1 scale with 1 meaning no species in common. Weighted UniFrac compared microbial ecosystems to one another based on the sequence distance in the phylogenetic tree and is based on the branch lengths of the phylogenetic tree that are weighted by relative abundances. Unweighted UniFrac compared microbial ecosystems to one another based on the sequence distance in the phylogenetic tree and is based on the sequence distance alone.

Differences were observed between the Bray Curtis Dissimilarity between the ceca and ileum (P = 0.001; Q = 0.001; **Figure 2A**). Additionally, the Bray Curtis Dissimilarity was different between all treatment groups (P = 0.001; Q ˂ 0.05; **Figure 2A**). Similar to Bray Curtis Dissimilarity, the Jaccard Diversity index was significantly different between that of the ceca and ileum (P = 0.001; Q = 0.001); **Figure 2B**). As well, the Jaccard Diversity index of the treatment groups was significantly different from one another (P = 0.001; Q ˂ 0.05; **Figure 2B**.

**Figure 2.**
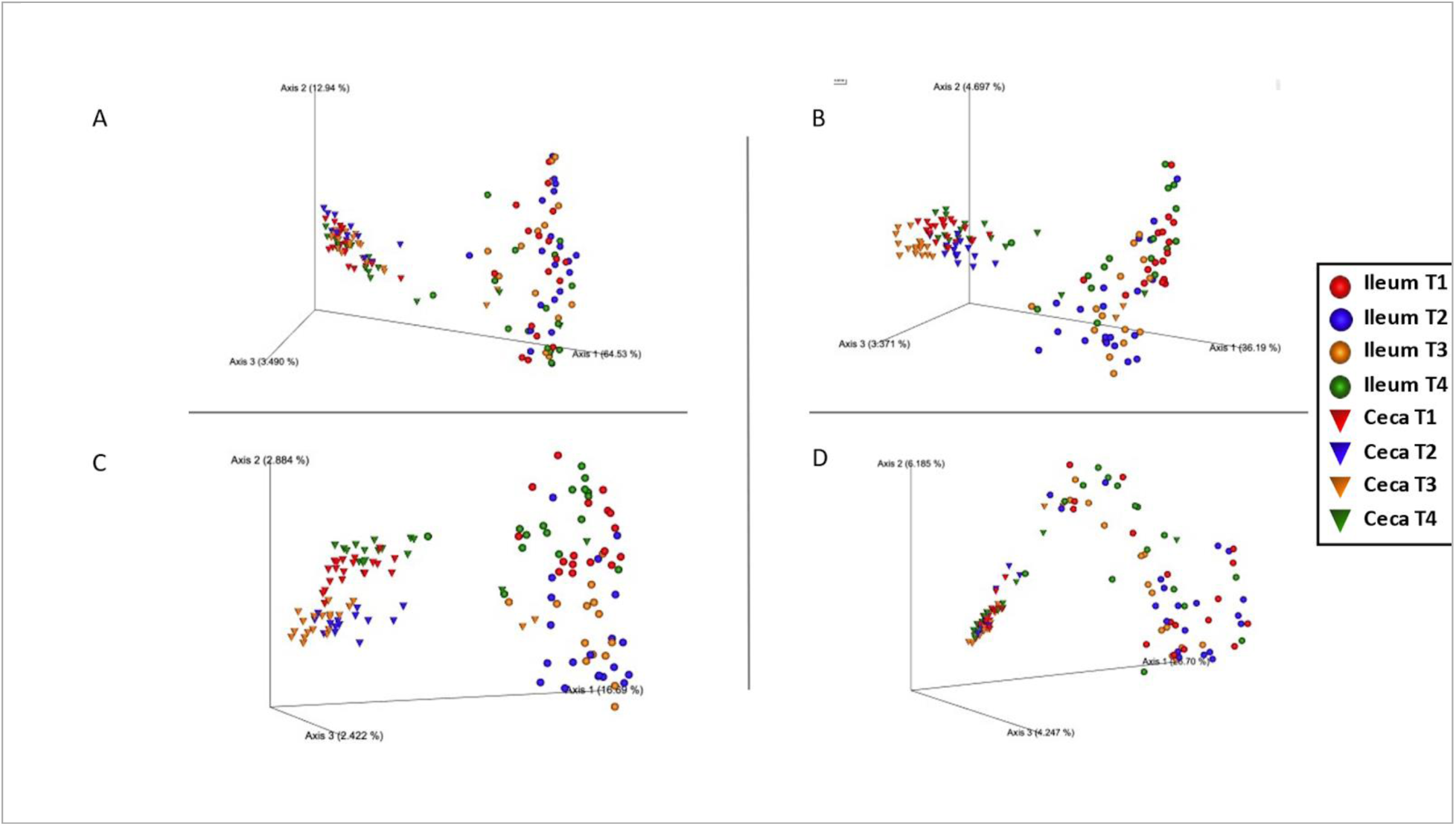
Beta Diversity of Ceca and Ileum. ^1^Treatment groups were as follows: 1). 2016 hen on 1940 diet; 2). 2016 hen on 2016 diet; 3). 1940 hen on 1940 diet; and 4). 1940 hen on 2016 diet *Beta Diversity of ceca and ileum from all treatment groups. A). Bray Curtis B). Jaccard distance C). Unweighted UniFrac D). Weighted UniFrac

The Operational Taxonomic Unit (OTU) distribution among hens from all treatment groups of the ileum and ceca is illustrated in **Figures 3** and **4**. OTU distribution at the phylum level among treatment groups of the ileum outlined the abundance of microbial communities which had no significant differences but illustrated Firmicutes as having the highest abundance in all treatment groups (**Figure 3A**). OTU distribution at the phylum level among treatment groups of the ceca outlined the abundance of microbial communities which had no significant differences but illustrated Bacteroidetes as having the highest abundance in all treatment groups (**Figure 3B**). OTU distribution at the genus level among treatment groups of the ileum outlined the abundance of microbial communities which had no significant differences, the top ten are illustrated which showed the highest abundance (**Figure 4A**). OTU distribution at the genus level among treatment groups of the ceca outlined the abundance of microbial communities which had no significant differences, the top ten are illustrated which showed the highest abundance (**Figure 4B**).

**Figure 3.**
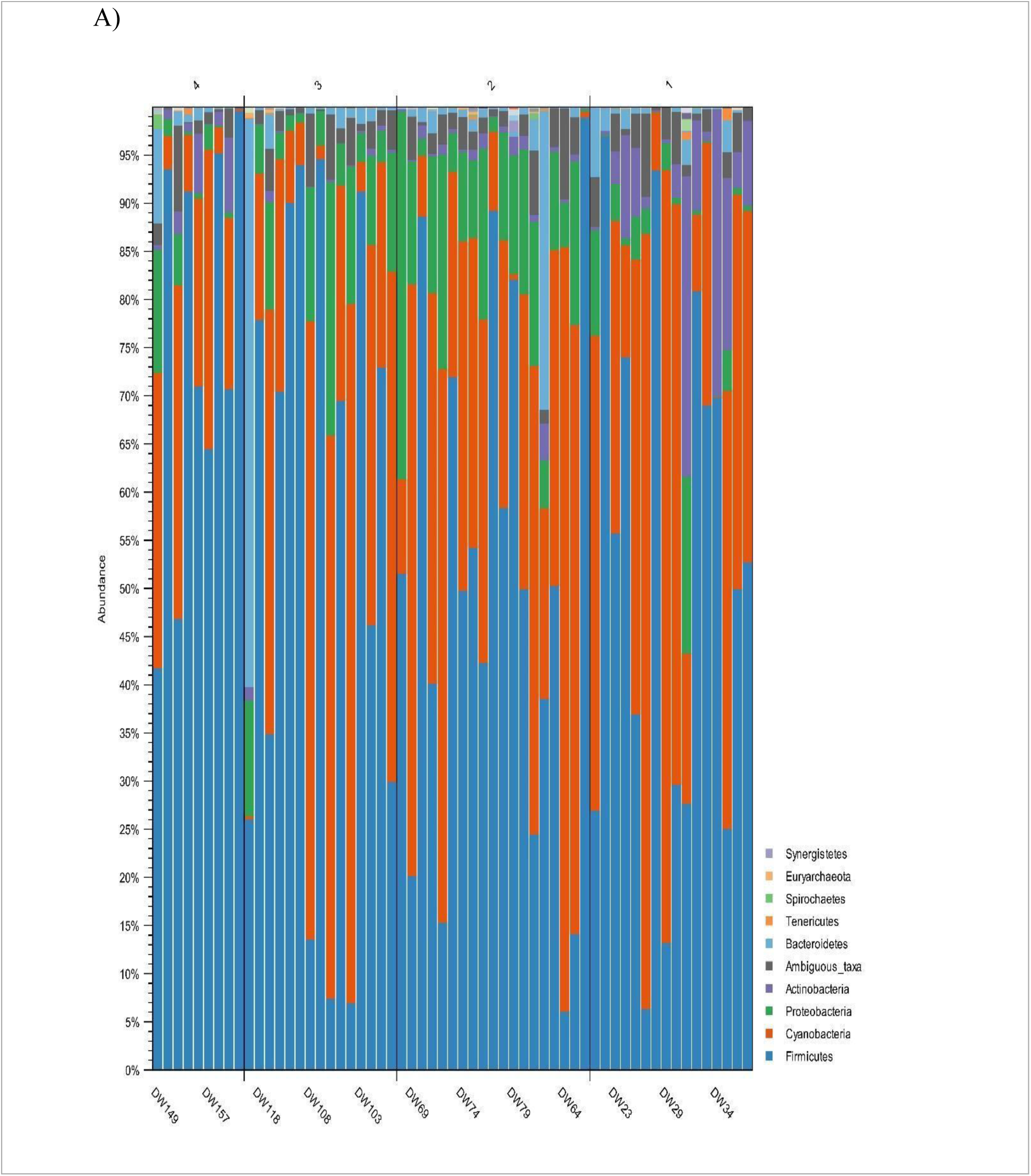
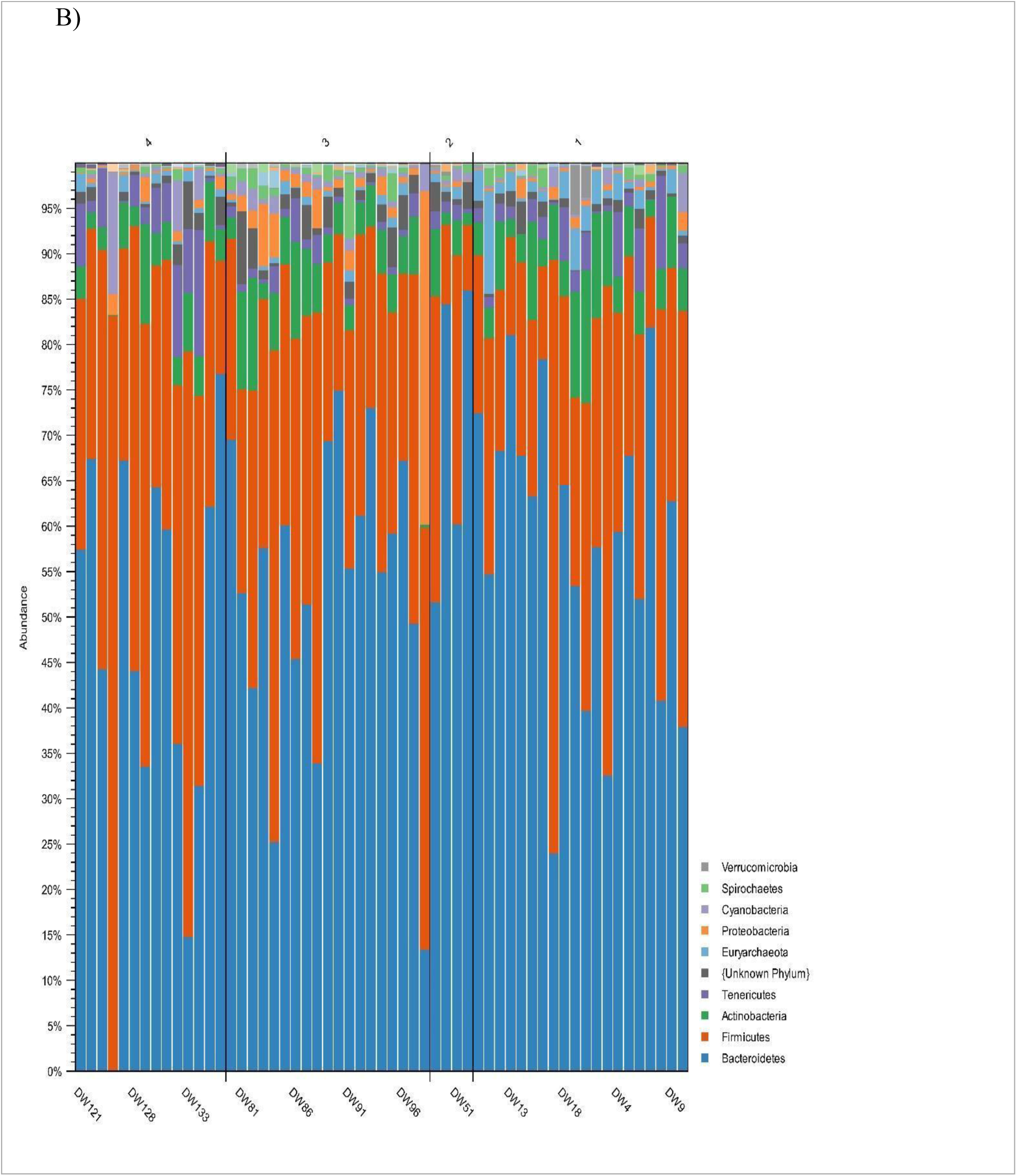
OTU Distribution at Phylum Level of Ileum and Ceca. ^1^Treatment groups were as follows: 1). 2016 hen on 1940 diet; 2). 2016 hen on 2016 diet; 3). 1940 hen on 1940 diet; and 4). 1940 hen on 2016 diet <u>^2^</u>Diagram A represents OTU distribution of phylum within the ileum <u>^3^</u>Diagram B represents OTU distribution phylum within the ceca

**Figure 4.**
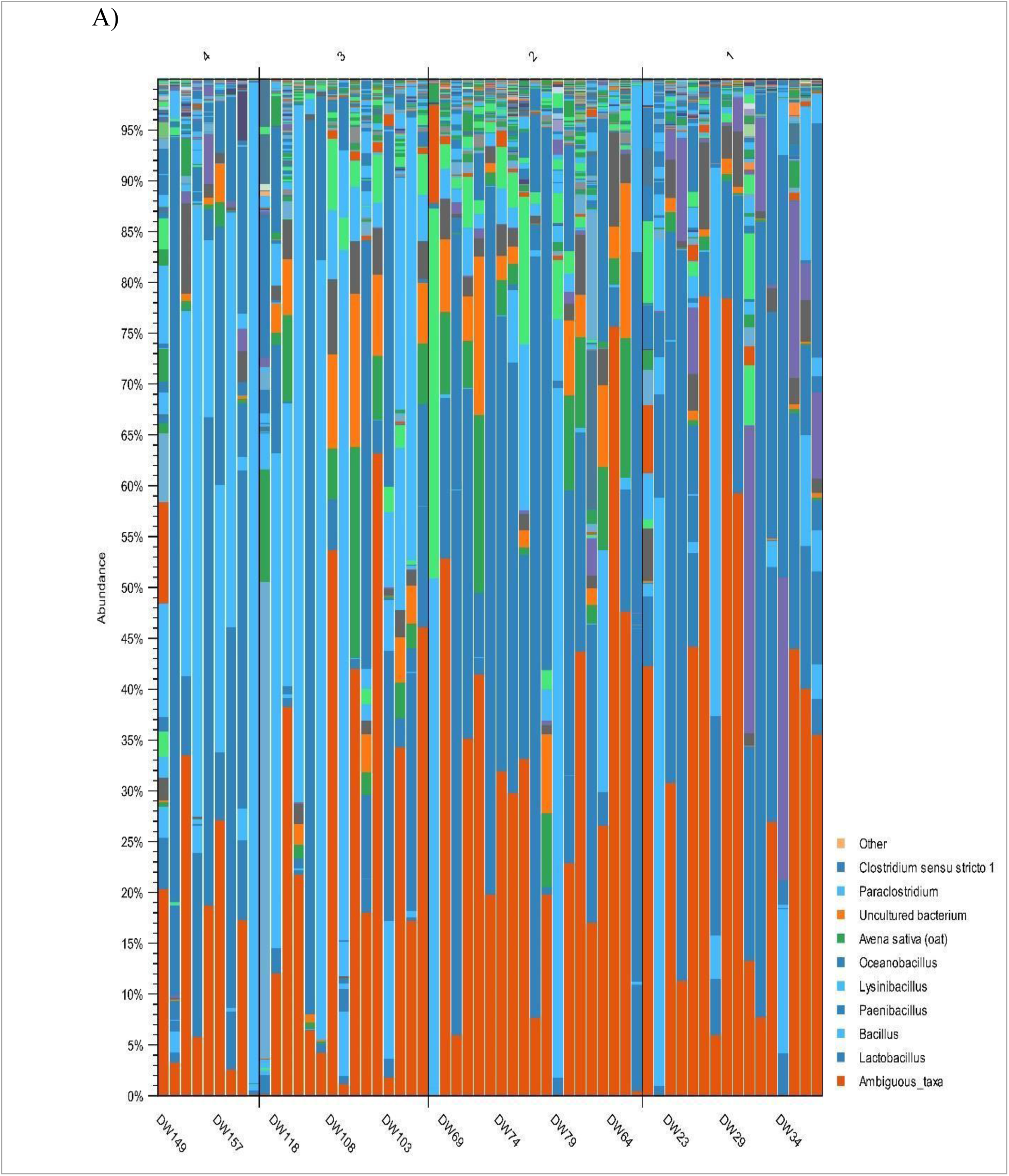
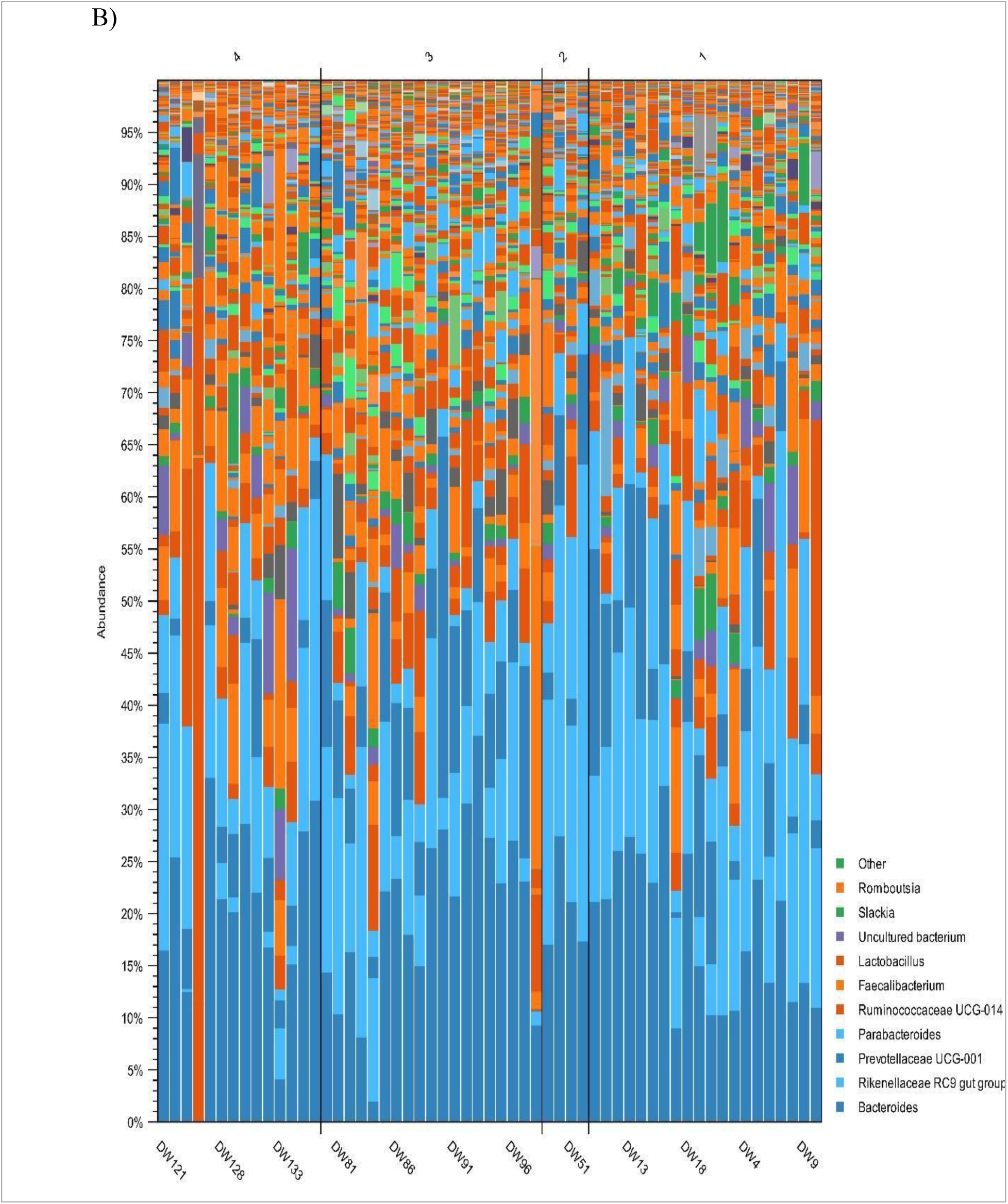
OTU Distribution at Genus Level of Ileum and Ceca. ^1^Treatment groups were as follows: 1). 2016 hen on 1940 diet; 2). 2016 hen on 2016 diet; 3). 1940 hen on 1940 diet; and 4). 1940 hen on 2016 diet <u>^2^</u>Diagram A represents OTU distribution of genus within the ileum <u>^3^</u>Diagram B represents OTU distribution genus within the ceca

### ANCOM

ANCOM, known as analysis of the composition of microbiomes, detected the differences in the microbial mean taxa abundance to provide exact differences when compared to beta diversity. In the ileum for ANCOM metrics, *Proteobacteria* had higher relative abundance in hens from treatment groups 2 and 3, while *Actinobateriota* had higher relative abundance in hens from treatment groups 1 and 4 at the Phyla level (**Figure 5A**). At the Genera level, hens from treatment groups 1 and 4 had higher relative abundance in *Aeriscardovia*, while hens from treatment groups 2 and 3 had higher relative abundance in both *Pseudomonas* and *Leuconostoc* (**Figure 5B**). In the ceca for ANCOM metrics, *Firmicutes* had the highest relative abundance across all treatment groups at the Phyla level (**Figure 6A**). At the Genera level, *Alloprevotella* had the highest relative abundance in hens from treatment groups 1 and 2, while *Leuconostoc* had the highest relative abundance in hens from treatment group 4, and *Pseudomonas* in hens from treatment group 4 (**Figure 6B**).

**Figure 5.**
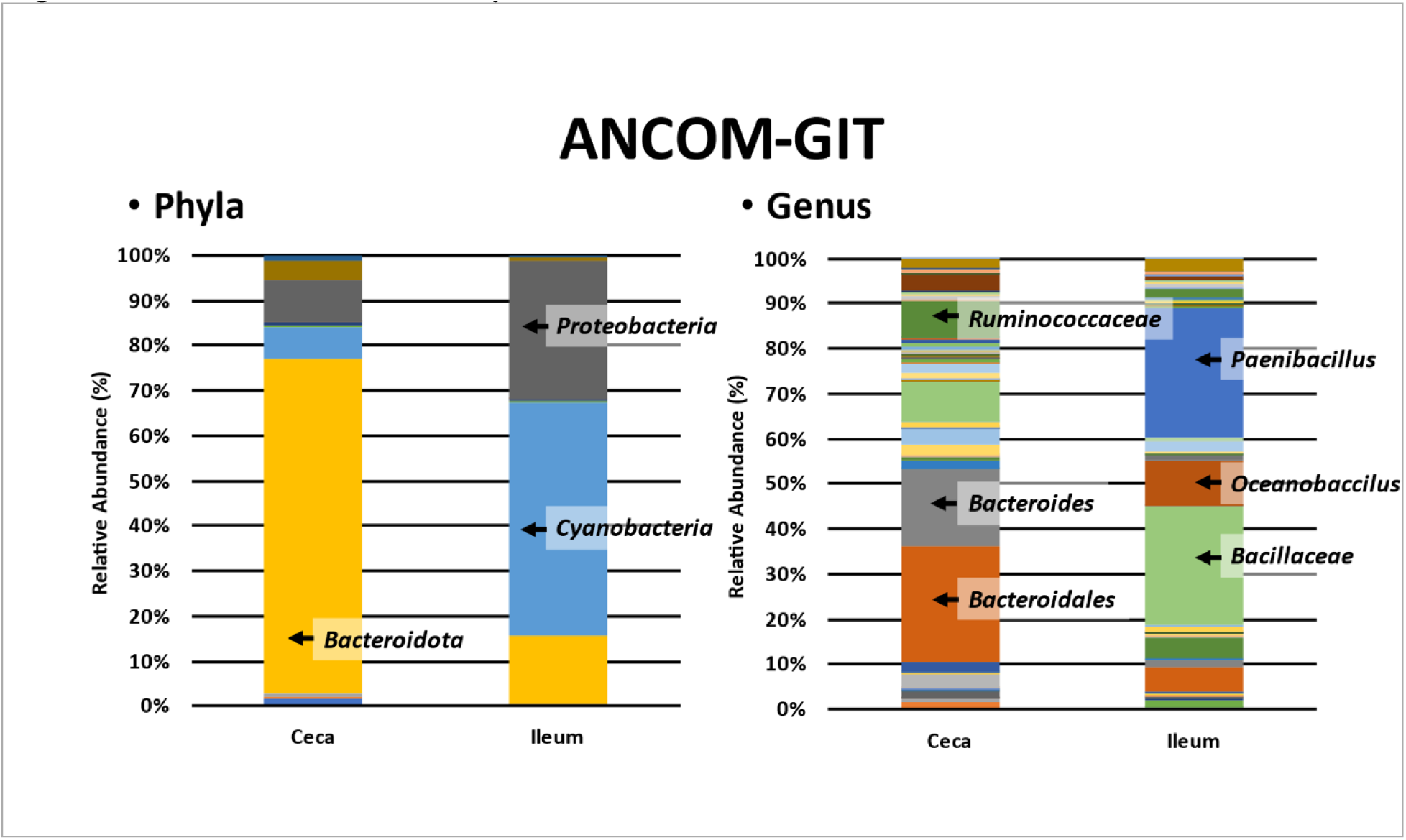
ANCOM of GIT at Phyla and Genus Levels. ^1^Relative abundance (%) of GIT

**Figure 6.**
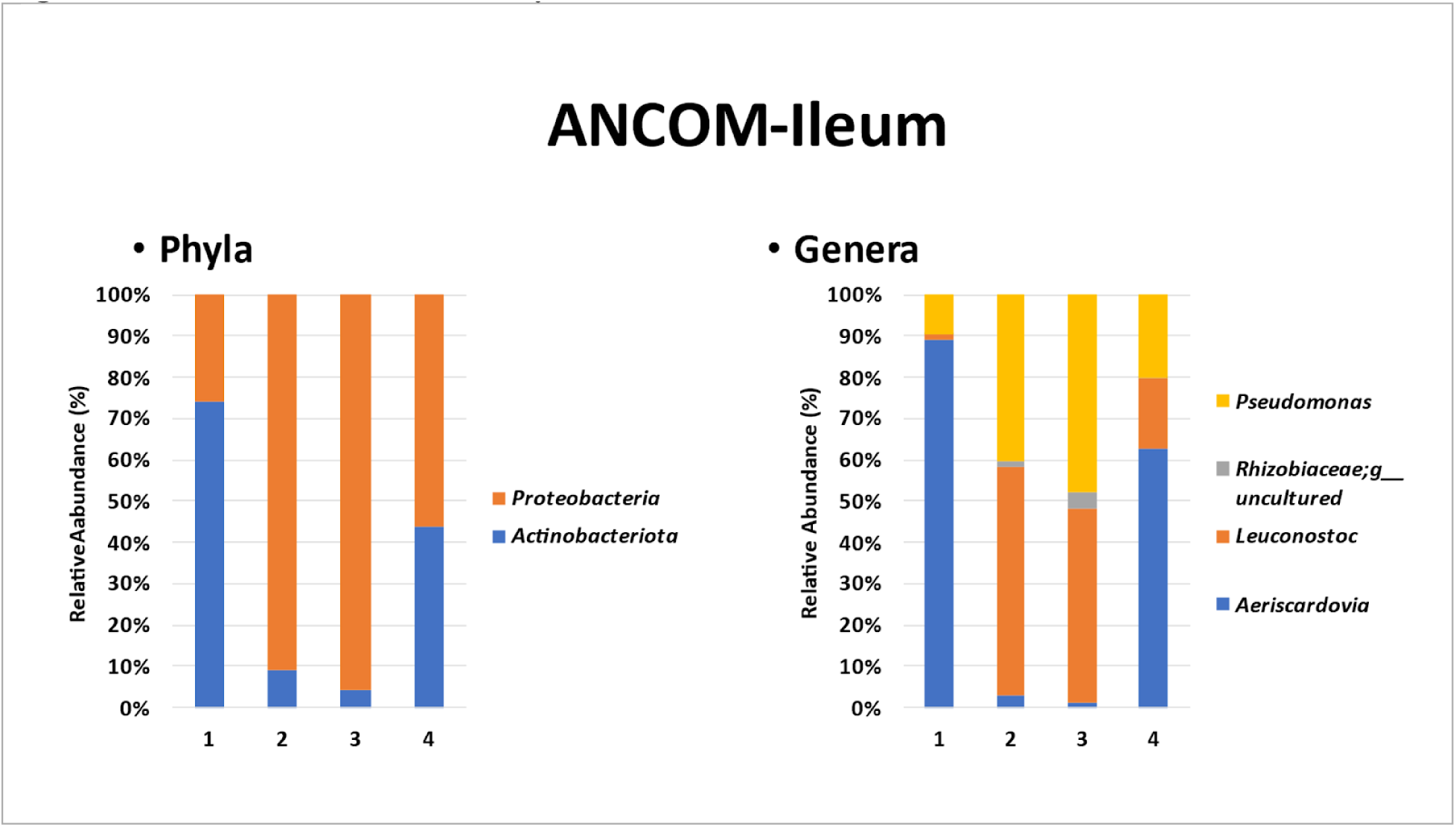
ANCOM of Ileum at Phyla and Genera Levels. ^1^Relative abundance (%) of ileum ^2^ANCOM at the phyla level of the ileum in **6A** ^3^ANCOM at the genera level of ileum in **6B**

**Figure 7.**
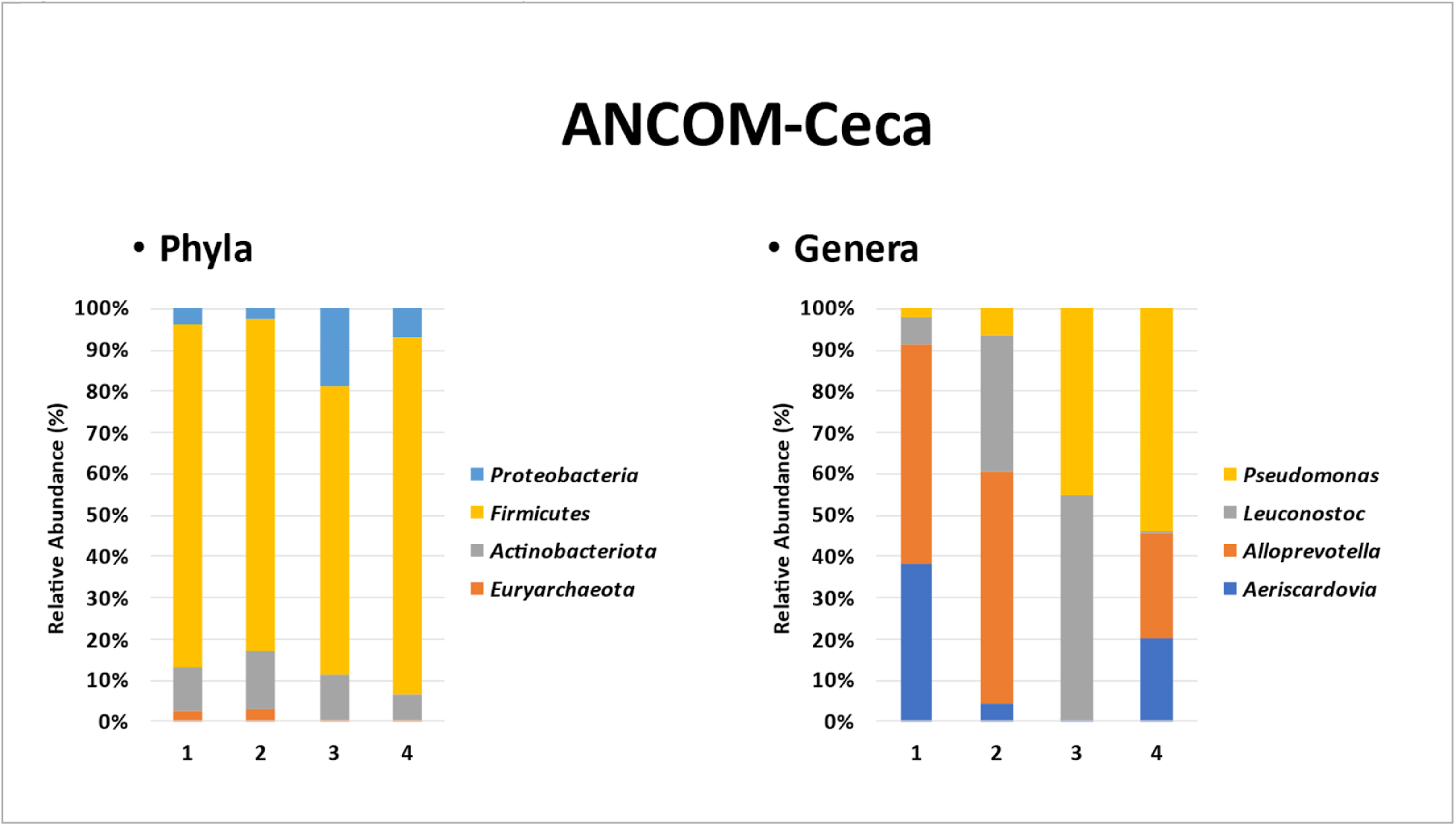
ANCOM of Ceca at Phyla and Genera Levels. ^1^Relative abundance (%) of ceca ^2^ANCOM at the phyla level of the ileum in **7A** ^3^ANCOM at the genera level of ileum in **7B**

## Discussion

A healthy gut has been deemed as an indicator of a healthy host, which in turn digests nutrients more efficiently for optimal production. The balance of microflora is the most important characteristic of a healthy gut. According to Lu et al., (2003) low bacterial diversity may be associated with reduced immunological and gut protective functions, as a rich and diverse bacterial population usually correlates with positive health status (Lu J, et al., 2003). The intestinal microbiota of chickens is affected by various factors and has been widely studied in conjunction with breed and host genotypes both having the ability to influence gut microbiota communities (Kers, J. G., et al., 2018). According to Mi et al., (2017), the host genotype has the greatest effect on the diversity of microbiota communities as well as the genotype of the host (Elokil AA, et al., 2020). Another contributing factor responsible for shaping the composition and function encoded by the microbiota in the gastrointestinal tract is diet. The efficiency of feed utilization is generally considered stable; however, it varies considerably among individuals fed identical diets and reared under the same conditions. The compounds within diets are important, providing growth substrates or microbes. In this study, ileum, and ceca microbiota composition was evaluated among two dissimilar leghorn strains that were fed on two different representative diets. Changes in the community composition, diversity, and richness of microbial communities in the ileum and ceca within the Alpha. Beta and ANCOM were identified. Factors such as diet and feeding regime could affect the population structure of chicken microbiota within host species (Huang, C. B et al., 2019).

### Alpha Diversity

The compartments of the digestive tract differ from one another, both morphologically and functionally, and have previously shown that their microbial composition is similarly distinct. Alpha diversity indices are used to estimate the diversity of microbial communities and it is believed that the richer the alpha diversity and species composition in microbiota, the better stability within the gut micro-ecosystems. Alpha diversity in this study measured the variation in the structure of the microbial community within individual samples of the ileum and ceca. Alpha diversity has been strongly correlated with variations in a specific gene function suggesting that gut microbiota with high diversity could potentially be more stable or labeled healthier than those with low diversity. The non-significance in alpha diversity among treatment groups measured within the ileum and ceca illustrated that the microbial communities were more evenly distributed. The low richness and evenness of some suggest that there were low numbers of species in both ileum and ceca consisting of a few dominating taxa. The microbiota of the small intestine, the section of the ileum as described in this study, could positively contribute to nutrient digestion and absorption processes, more specifically water and mineral absorption. Microbial compositions of both the environment and gastrointestinal tract vary considerably in poultry production and can be affected by factors such as age and production batch, therefore, we tried to limit many factors such as age, breed, gender, and house to minimize the fluctuation that exists within the gut microbiome. Results indicate that alpha diversity measured in this study can contribute to functional inference involving the underlying microbiota mechanism. The lack of differences can be attributed to the hens being exposed to the same environmental sources of microorganisms that are known to have an effect on overall diversity as well as the richness of the microbiome in poultry.

### Beta Diversity

Different nutrient requirements may also affect the composition and abundance of microbiota in different intestinal segments. Beta diversity was used to compare differences in species diversity between multiple samples. Beta diversity in this study measured the similarity or dissimilarity of microbial communities between samples of the ileum and ceca; thus, making it a useful technique to observe changes within the community composition. Although beta diversity consisting of Bray-Curtis, Jaccard, and Unweighted UniFrac produced significant variations in the ileum and ceca, there was no significant effect among hens within treatment groups with Weighted UniFrac indicating that distinct phylogenetic groups are possible, but the prevalent abundance of taxonomic groups persist. Lower UniFrac distance indicates a more similar bacterial composition between the samples. The two most dominant bacterial phyla in chicken belong to Firmicutes and Bacteroidetes (Magne, F. et al., 2020). Firmicutes species are associated with the decomposition of polysaccharides and the production of butyrate (Flint, H. J et al., 2012). Bacteroidetes species degrade complex carbohydrates and synthesize propionate via the succinate pathway.

### ANCOM

ANCOM accounts for the underlying structure in the data and can be used for comparing the composition of microbiomes in two or more populations (Siddhartha Mandal et al., 2015). The microbiome compositional diversity did not group the samples at family and species levels according to the diet. The function of the ileum has a major role in water and mineral absorption with evidence of contributing to starch and fat digestion and is generally known to harbor a diverse and numerically more significant population of bacteria when compared to the other two intestine segments. The ileum microbiota at times can intermix with the microbiota of the ceca. Within the ileum, at the phyla level, Proteobacteria (gram-negative) had higher relative abundance followed by Actinobacteriota (gram-positive), both non-spore-forming bacteria. Hens belonging to treatment group 2 (2016 hen on 2016 diet) and treatment group 3 (1940 hen on 1940 diet) had an increased abundance of Proteobacteria suggesting that genetics had a greater influence on observed bacteria. In that same respect, similar results were seen in hens belonging to treatment group 1 (2016 hen on 1940 diet) and treatment group 4 (1940 hen on 2016 diet) having an increased abundance in Actinobacteria suggesting that the influence of diet may have been attributed to the observed presence of bacteria. Actinobacteria have a number of important functions, including the degradation/decomposition of all sorts of organic substances such as cellulose, polysaccharides, protein fats, organic acids, etc. The abundance of Actinobacteria in hens belonging to treatment groups 1 and 4 may be a direct result of the composition of the diets provided to these hens. The findings in this study were similar to the results from research conducted by Ngunjiri et al., (2019) and Wang et al., (2019) where the majority of phyla belonged to both Proteobacteria and Actinobacteria of the ileum (Wiersema, M. L et at., 2021). At the genera level, hens belonging to treatment groups 1 (2016 hen on 1940 diet) and 4 (1940 hen on 2016 diet) had a dominant abundance of Aeriscordovia. Hens belonging to treatment groups 2 (2016 hen on 2016 diet) and 3 (1940 hen on 1940 diet) had a dominant abundance of Leuconostoc.

A complex bacterial community is found in the ceca primarily due to the longer digestive transit times making it an important site for the recycling of urea, water regulation, and carbohydrate fermentations while contributing to intestinal health and nutrition. The abundance of Firmicutes, the major bacterial phyla observed, was dominated in all treatment groups within the ceca which were similar to research conducted by Hamid et al., (2019). Proteobacteria and Actinobacteria were the second and third abundant components observed at the phyla level in the ceca which typically accounts for 2-3% of total microbiota. Despite variations observed, representatives of the phyla mentioned above are found in the ceca of nearly all adult chickens.

Hens belonging to treatment groups 1 and 2 of the 2016 strain were more abundant in Alloprevotella (a fiber degradation bacteria) at the genera level and hens belonging to treatment groups 3 and 4 of the 1940 strain were more abundant in Pseudomonas suggesting that genetics had a greater influence on bacteria observed rather than diet, however, diet may have attributed to the differences of bacteria amounts found. The variations validated by the ANCOM results suggest that there were some bacteria that persevered genetically throughout the treatment groups despite the differences in the nutritional composition of the diets provided. Direct comparison of the microbiota composition established between laying hen strains has the ability to provide valuable insight into the association between microbiota, genetic and nutritional composition (James J. Kozich et al., 2013).

## Conclusion

The chicken gut microbiota is colonized with complex microbial communities, known to play an important role in overall health and performance. Distinct microbiota between the ileum and ceca were observed for both alpha and beta diversity and ANCOM. Results from this study suggest that genetic makeup in conjunction with the nutritional composition of laying hens has the ability to influence both ileal and cecal microbiota. The comparison of microbiota composition between laying hen breeds can provide valuable insight into the linkage between microbiota distribution and genetic architecture. This information can collectively be beneficial in explaining the behavior of the intestinal ecosystem in order to proactively modulate necessary changes needed to enhance overall health status. Furthermore, it remains to be determined whether the interplay of diet and genetics singlehandedly fosters the changes observed within the microbiome of the gastrointestinal tract, however, the results from this study provide a vital step in identifying the roles exhibited by the various bacteria mentioned above.

